# Elucidating Molecular Mechanisms Governing TNF-Alpha-Mediated Regulation of Amyloid Beta 42 Uptake in Blood-Brain Barrier Endothelial Cells

**DOI:** 10.1101/2025.01.28.635286

**Authors:** Vrishali S. Salian, Geoffry L. Curan, Val J. Lowe, Xiaojia Tang, Krishna R. Kalari, Karunya K. Kandimalla

## Abstract

Cerebrovascular inflammation is prevalent in a majority of Alzheimer’s patients. Inflammatory cytokines, such as tumor necrosis factor-alpha (TNF-alpha), circulating in the plasma have been shown to cause the inflammation of blood-brain barrier (BBB) endothelium lining the cerebral microvasculature. The BBB inflammation has been implicated in the increase of toxic Aβ accumulation within Alzheimer’s disease (AD) brain. TNF-alpha in the peripheral circulation can aggravate the accumulation of amyloid-beta (Aβ) peptides in Alzheimer’s disease brain. In the current study, we have shown that the exposure to TNF-alpha leads to an increase in Aβ42 accumulation in mice and BBB endothelial cells in vitro. Moreover, dynamic SPECT/CT imaging in wild-type (WT) mice infused with TNF-alpha increased the permeability and influx of Aβ42 into the mice brain. In addition, our results show that TNF-alpha modifies the expression of cofilin, actin, and dynamin, which are critical components for Aβ endocytosis by BBB endothelial cells. These results offer a mechanistic understanding of how TNF-alpha may promote Aβ accumulation at the BBB and the underlying interactions between inflammation and Aβ exposure that drives BBB dysfunction. Hence, a therapeutic intervention aimed at addressing cerebrovascular inflammation in Alzheimer’s disease may potentially reduce Aβ induced cerebrovascular toxicity in Alzheimer’s disease brain.

**Significance statement:** Increased levels of TNF-alpha circulating in the plasma are considered significant factors in the consequences of Aβ pathology in Alzheimer’s disease, where it can promote cerebrovascular inflammation and BBB dysfunction. However, the role of TNF-alpha, in exacerbating Aβ pathology by increasing Aβ accumulation at the BBB endothelial cells remains only partially understood. In this study, we demonstrated that TNF-alpha enhances Aβ42 accumulation in the BBB endothelium by altering the expression of the BBB endocytosis machinery, specifically cofilin, actin, and dynamin. These findings are anticipated to contribute to the development of therapeutic approaches aimed at addressing elevated cytokine levels in Alzheimer’s disease.

**Graphical Abstract.**
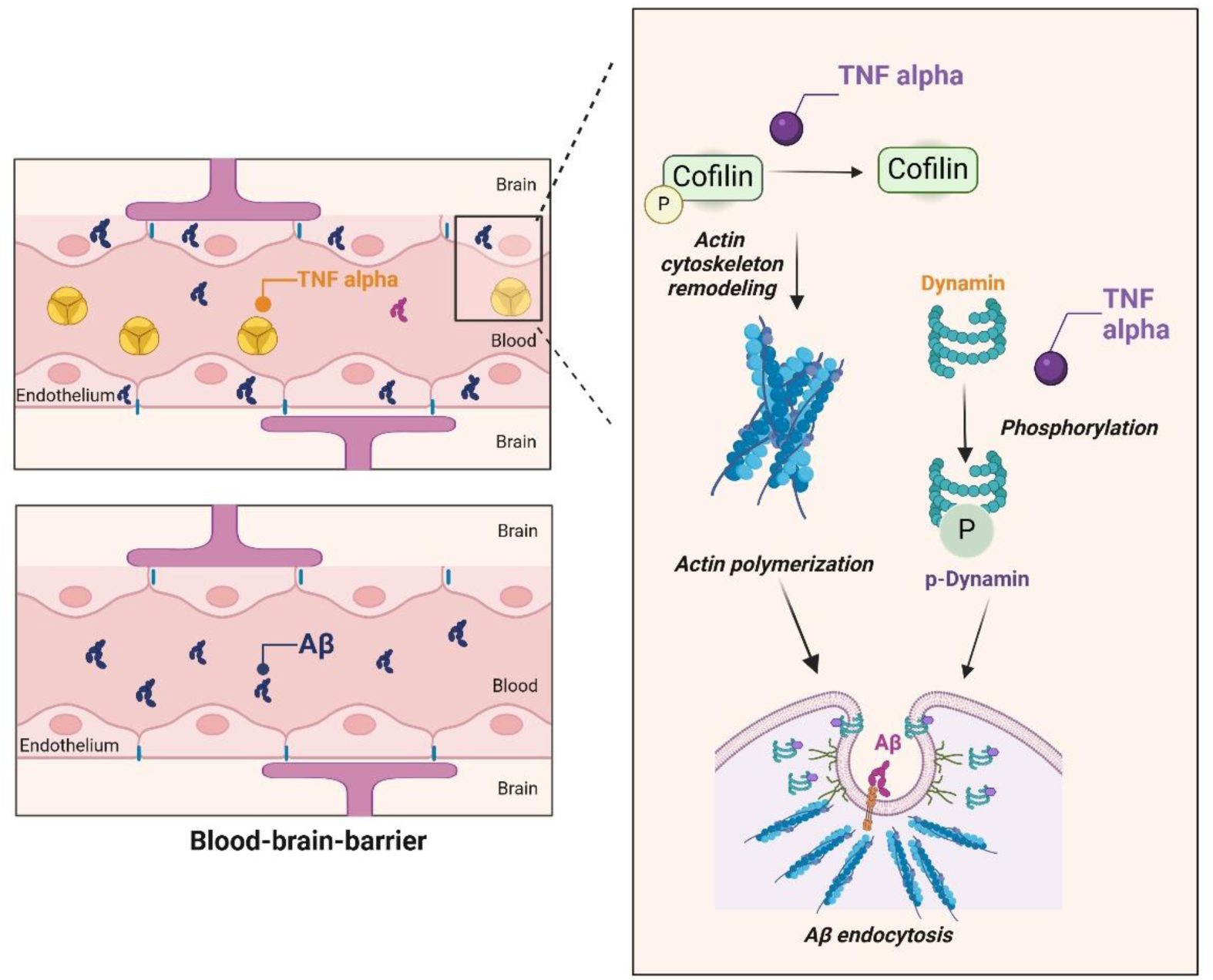

## Introduction

Alzheimer’s disease is a neurodegenerative disease characterized by the accumulation of amyloid beta (Aβ) peptides in the brain parenchyma and within cerebral vasculature(Hippius and Neundorfer 2003; Selkoe 2001). The endothelial cells lining the cerebrovascular lumen, commonly known as the blood-brain barrier (BBB), regulates dynamic equilibrium of Aβ levels between the brain and plasma (Wang et al. 2021). The luminal-to-abluminal transport of Aβ is mediated by the receptor for advanced glycation end products (RAGE)(Deane et al. 2003), while abluminal-to-luminal transport is handled by low-density lipoprotein receptor-related protein-1 (LRP- 1) and efflux transporters such as P-glycoprotein (Shibata et al. 2000; Storck et al. 2016; Park et al. 2014).

Once Aβ circulating in the blood is endocytosed by the RAGE receptor, transcytosis machinery within the BBB endothelium regulates its transcytosis to the brain parenchyma. Cofilin, actin filaments, and dynamin are some of the key components driving endothelial endocytosis. Cofilin is an actin-binding protein that severs actin filaments, promoting invagination of the endothelial plasma membrane(Alhadidi, Bin Sayeed, and Shah 2016, 2017) which facilitates the formation of endocytic vesicles. Dynamin, a cell surface GTPase protein, is involved in membrane fission by encircling the necks of budding endocytic vesicles(Antonny et al. 2016; Ferguson and De Camilli 2012; Schmid and Frolov 2011). Although, endocytosis is known to be disrupted in Alzheimer’s disease(Kommaddi et al. 2018; Wang et al. 2023), the role of cofilin and dynamin expression changes on the dysregulation of Aβ endocytosis by the BBB endothelial cells remain elusive and poorly understood.

Chronic inflammation in the brain plays a pivotal role in AD progression. Pre-existing Aβ pathology may render the BBB more sensitive to proinflammatory cytokines(Amtul et al. 2019), which promotes BBB dysfunction. Tumor necrosis factor (TNF) alpha is a proinflammatory cytokine that is upregulated in the plasma of AD patients compared to healthy controls and individuals with mild cognitive impairment(Khemka et al. 2014; Kim, Lee, and Kim 2017; Brosseron et al. 2014). Moreover, it causes BBB dysfunction and accelerates Alzheimer’s disease progression(Gao and Bayraktutan 2023; Nishioku et al. 2010). TNF-alpha potentially promotes BBB dysfunction by stimulating actin cytoskeleton reorganization via filamentous actin disruption and cytoskeleton contraction, which promotes transendothelial flux via the formation of a widened intracellular space(Kumar et al. 2009; Bogatcheva et al. 2003). Thus, TNF-alpha could facilitate Aβ accumulation in endothelial cells by affecting the expression of intracellular trafficking machinery components, such as cofilin, actin, and dynamin.

Although these findings provide circumstantial evidence of a possible crosstalk between inflammation and Aβ pathology, this link has not been systematically examined, and the underlying mechanisms remain elusive. In the present study, we focused on the influence of TNF-alpha on the accumulation of Aβ42 in BBB endothelial cells and changes in endocytosis machinery that drive this phenomenon.

## Methods

### Animals

Female wild-type mice (B6SJLF1) were procured from Jackson Laboratory (Bar Harbor, ME) and housed in a virus-free barrier facility with 12-hour light-dark cycles. They were provided ad libitum with pellet food and purified water. The animal studies described in this study were conducted in accordance with the National Institutes of Health Guide for the Care and Use of Laboratory Animals. The Institutional Animal Care and Use Committee at Mayo Clinic, Rochester, MN, approved the study protocols, which were recorded as per the ARRIVE reporting guidelines.

### Cell culture

The immortalized human cerebral microvascular endothelial cell (hCMEC/D3) line was generously provided by P-O Couraud of Institut Cochin, France. The endothelial cells were cultured according to the protocols provided by the Couraud group(Daniels et al. 2013). Briefly, the hCMEC/D3 cells were cultured in endothelial basal media (Sigma-Aldrich, St. Louis, MO), supplemented with various reagents that support cell growth. These included a chemically defined lipid concentrate (1% v/v, ThermoFisher Scientific, Waltham, MA), penicillin-streptomycin (1% v/v, Sigma-Aldrich, St. Louis, MO), recombinant human fibroblast growth factor (1 ng/mL, PeproTech, Rocky Hill, NJ), hydrocortisone (1.4 μM, Sigma-Aldrich, St. Louis, MO), ascorbic acid (5 μg/mL, Sigma-Aldrich), HEPES buffer (10 mM), and fetal bovine serum (5% or 1%), Atlanta Biologicals, Flowery Branch, GA). This complete medium, along with the supplements, is referred to as D3 media. The polarized hCMEC/D3 monolayers were cultured in collagen-coated (Corning, MA) 35 mm glass coverslip bottomed dishes (Mattek, MA) or 6-well plates (Corning Costar, MA) under 5% CO_2_ at 37°C. Cells from passages 32–34 were used for the experiments. The medium was changed every 2 days until reaching confluence (∼4 days total). These monolayers were initially cultured in 5% serum-containing D3 media.

### Isolation and culture of primary porcine brain microvascular endothelial cells

Primary porcine brain microvascular endothelial cells (PBMECs) were isolated using the method described by(Gali et al. 2019). The PBMECs enriched from the porcine brain microvasculature were seeded in t-150 flasks pre-coated with 150 μg/ml type-I rat tail collagen and maintained at 37 °C under 5% CO_2_ in Earle’s medium (M199 1X). This medium was supplemented with 1% (v/v) penicillin, 1% (v/v) streptomycin, 1% (w/v) gentamycin, 1 mM L-glutamine, 10% (v/v) horse serum. The PBMEC’s were treated for 24 h with puromycin dihydrochloride (4 μg/ml) to kill pericytes, and maintained in growth medium for 5 days, to allow for the proliferation of PBMECs. Subsequently, the PBMECs were sub-cultured into 6-well plates for flow cytometry studies.

### Dynamin mRNA gene expression in APP/PS1 transgenic mice

The mRNA gene expression patterns of dynamin were analyzed using data from APP/PS1 transgenic (TG) mice as well as wild-type (WT) counterparts, all at 8 months of age obtained from the gene expression omnibus (GEO) database under accession number GSE85162. Gene expression analysis was conducted using the Affymetrix HT MG-430 PM array, focusing on the hippocampus and prefrontal cortex regions of both male and female APP/PS1 transgenic mice.

### Aβ peptide film preparation and reconstitution

The unlabeled and fluorescein isothiocyanate (FITC) conjugated Aβ peptides were obtained from Aapptec (Louisville, KY) at 95% purity, as determined by high-performance liquid chromatography-mass spectrometry. The monomeric solutions of Aβ peptides were prepared in accordance with previously published methodologies.(Chromy et al. 2003)

### Radioiodination of Aβ peptides

The Aβ42 peptides (Aapptec, Louisville, KY) were labeled with the ^125^I radionuclide (PerkinElmer Life and Analytical Sciences, Boston, MA) utilizing the chloramine-T procedure as previously described(Kandimalla et al. 2005; Poduslo, Curran, and Berg 1994). The radio-iodinated proteins were separated from the free ^125^I through dialysis against 0.01 M Dulbecco’s phosphate-buffered saline (DPBS) at pH 7.4 (Sigma-Aldrich, St. Louis, MO). The integrity of the radio-iodinated Aβ42 (^125^I-Aβ42) was assessed using the trichloroacetic acid (TCA) precipitation method(Kandimalla et al. 2005). The radio-iodinated proteins were separated from the free ^125^I through dialysis against 0.01 M DPBS at pH 7.4 (Sigma-Aldrich, St. Louis, MO). The integrity of the radio-iodinated Aβ42 (^125^I-Aβ42) was assessed using the TCA precipitation method

### Plasma pharmacokinetics of ^125^I-Aβ42 peptides in wild-type mice following TNF - alpha infusion

This experiment was conducted in accordance with the methodologies delineated in our previous publication. Briefly, the internal carotid artery and femoral vein of female wild-type (WT) mice (4 months of age) were catheterized under general anesthesia (1.5% isoflurane; 4L/min oxygen). The TNF-alpha (25 μg/kg) was infused at a rate of 5 μL/min via the internal carotid artery and allowed to circulate for 1 hour. A single dose (150 μCi) of ^125^I-Aβ42 in phosphate-buffered saline (PBS) (100 μL) was bolus injected via the femoral vein. Then, 20 μL blood samples were withdrawn and collected in heparinized capillary tubes at 0.25, 1, 3, 5, 7, 10, 15, 30, and 40 minutes following the IV bolus injection. The collected blood samples were diluted with saline and centrifuged to separate the plasma. Intact ^125^I-Aβ42 in the plasma was assayed following TCA precipitation in a gamma counter (Cobra II; Amersham Biosciences, Piscataway, NJ). At the end of the experiment, the mice were euthanized by transcardial perfusion with excess PBS to flush the remaining radioactivity from the vasculature. Subsequently, the brain was removed from the cranial cavity and dissected into various anatomical regions (cortex, caudate putamen, hippocampus, thalamus, brain stem, and cerebellum). Individual brain regions were assayed for ^125^I activity using a two-channel gamma counter. The ^125^I-Aβ42 plasma concentration vs time profile was fitted to a two-compartment model using SAAMII software (Nanomath LLC). Permeability of ^125^I-Aβ42 at the BBB in each brain region was assessed as the permeability–surface area (PS) product (milliliter/gram/second), determined using the equation:

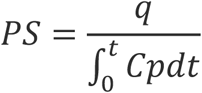

Where q is the extravascular amount of ^125^I-Aβ42 per gram of tissue (μCi/g) and 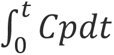 is the area under the plasma concentration versus time profile (minute × microcurie per milliliter) from 0 to 40 minutes, calculated using the linear-trapezoidal method. The % of the injected dose that permeated into the whole brain per gram tissue (% ID/g) was estimated using the equation:

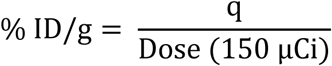

### Dynamic SPECT/CT imaging of ^125^I-Aβ42 influx to the brain

These experiments were also performed as described in our earlier publication(Swaminathan et al. 2018). Briefly, WT mice (*n* = 4 mice per group) were anesthetized, femoral vein as well as carotid artery were catheterized. TNF-alpha (25 μg/kg) or PBS control was infused via the internal carotid artery. One hour after the end of infusion, a bolus of ^125^I-Aβ42 (500 μCi) was injected into the femoral vein. The mice were then immediately placed in a SPECT/CT scanner and imaged at 1 min intervals over the course of 40 min, followed by a 5 min CT scan.

The blood-to-brain influx of ^125^I-Aβ42 was determined by Gjedde-Patlak analysis(Patlak, Blasberg, and Fenstermacher 1983), and involves plotting the following;

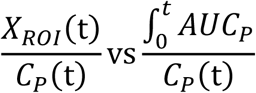

Where:

- X_ROI_(t) is the amount of ^125^I-Aβ42 activity (µCi) in the brain ROI at time t (min)
- C_p_(t) is the concentration in plasma (µCi /mL) at time t (min)
- 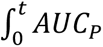 is the area under the plasma concentration vs time profile (µCi·min/mL) from time 0–t obtained by the logarithmic trapezoidal method. The plasma concentration at each imaging time point was predicted using the two-compartment pharmacokinetic parameters obtained in the preceding experiment. The brain influx, Ki (mL/min), is predicted from the slope obtained from the linear portion of the curve.

### Cofilin knockdown

Cofilin knockdown was performed in hCMEC/D3 monolayers after reaching 60%- 70% confluency. The knockdown procedure utilized a siRNA kit containing cofilin siRNA (Santa Cruz, TX), RNAi max (Invitrogen, CA), and Opti-MEM (Gibco, NY) with reduced serum. Following transfection, the cells were allowed to recover for 24 hours prior to the preparation of cell suspensions for flow cytometry, immunofluorescence staining for confocal microscopy, and western blot analysis.

### Phalloidin staining

Cells were grown on 35 mm glass-bottom coverslips. Following cofilin knockdown, the hCMEC/D3 monolayers were treated with TNF-alpha (20 ng/mL, 24hrs, 37C) in low serum media, washed with PBS to remove the treatment media. After fixing with 4% PFA for 15 minutes at room temperature, the cells were permeabilized with 0.1% Triton X-100 in PBS for 10 minutes. The permeabilized cells were incubated at 25°C for 1 h with AF-647 phalloidin (Abcam) in PBS containing 1% BSA to stain for F-actin. Subsequently, the cells were treated with 3 µM DAPI (Invitrogen, MA) for 10 minutes at room temperature and mounted using ProLong gold antifade mounting medium (Invitrogen, MA). The cells were imaged with Nikon Confocal A1Rsi NSIM confocal microscope using a 60X,1.4NA objective (University of Minnesota Imaging Center). Data was acquired using NIS Elements AR software (v.5.6, Nikon Instruments USA, Melville, NY). The images were quantified using Fiji software where the mean fluorescence intensity of each image was quantified and normalized to the cell number, determined using the cell counter function in Fiji.

### Preparation of cell lysates and western blotting

The hCMEC/D3 monolayers were treated with TNF-alpha (20 ng/mL, 24 hours, 37°C) (R&D systems, MN) or blank medium. Following the treatment, the cells were washed three times with ice-cold PBS and lysed using radioimmunoprecipitation assay (RIPA) buffer containing protease and phosphatase inhibitors (Sigma-Aldrich, St. Louis, MO). The total protein concentration of the whole-cell lysates was determined using the bicinchoninic acid assay (Pierce, Waltham, MA).

Whole-cell lysates containing 20-30 µg of protein were loaded on 6%-12% polyacrylamide gels, and resolved by SDS-PAGE under reducing conditions (Bio-Rad Laboratories, Hercules, CA). Then the proteins electrophoretically transferred to a nitrocellulose membrane (0.2 µm pore size) and were blocked for 1 hour at room temperature using 5% w/v milk protein in tris-buffered saline with 0.1% Tween20 detergent (TBST). Primary antibodies from Cell Signaling Technology (Danvers, MA) targeting p-dynamin, p-cofilin (serine 3), cofilin, and GAPDH (protein loading control) were used to probe respective proteins. After overnight incubation with the primary antibody (1:1000 dilution) at 4 °C, the nitrocellulose membrane was washed with TBST to remove any unbound antibody and incubated with a goat anti-rabbit near-infrared (800 nm) IR-dye-conjugated secondary antibody (Licor, NE) (1:2000) dilution for one hour at room temperature. The proteins on the nitrocellulose membrane were visualized using an Odyssey Licor instrument (Odyssey CLx; LI-COR Inc.) and quantified using Image Studio Lite 5.2 software. The intensity of the bands on the nitrocellulose membrane were normalized to the intensity of the GAPDH band, which served as the loading control.

### Flow cytometry

The polarized hCMEC/D3 monolayers or PBMECs were treated with TNF-alpha (10/20 ng/mL, 24 hours, 37°C) or blank medium followed by incubation with FITC-Aβ42 (1 μM) peptide for 60 minutes. Similar experiment was also conducted by incubating the hCMEC/D3 monolayers with 100 nM Alexa fluor-647 (AF-647) labeled insulin (Nanocs, New York, NY) for 20 minutes at 37°C. The cell monolayers were washed thoroughly with PBS, the cells were dislodged with trypsin-EDTA and fixed with 4% paraformaldehyde. Then the intracellular fluorescence of FITC-Aβ42 or AF-647 insulin was quantified using a BD FACSCalibur flow cytometer equipped with 488 nm (FITC- Aβ42) and 633 nm (AF-647 insulin) lasers, respectively. The data was acquired using BD CellQuest Pro and analyzed using FlowJo software. The uptake of FITC-Aβ42 or AF-647 insulin was presented as histograms of intracellular fluorescence and the median fluorescence intensities were plotted as bar charts.

### Immunocytochemistry and Cellular Imaging

Polarized hCMEC/D3 cell monolayers were cultured on 35 mm glass-bottom dishes in D3 media, which was then changed to low serum D3 medium (1% FBS) one day prior to the experiment. The cells were stratified into four treatment groups: control, TNF-alpha, dynasore, a dynamin inhibitor, and a combination of TNF-alpha and dynasore. The cells were maintained at 37°C and treatment conditions are described in Table 1. After the treatment period, the cells were exposed to FITC-Aβ42 (1 µM) for 30 minutes, followed by co-incubation with Lysotracker red (75 nM) (Invitrogen-Molecular Probes, CA) for another 30 minutes. The cells were then washed with PBS, fresh D3 media was added, and the cells were Imaged live with confocal microscopy.

**Table 1.**

| **Group** | **Pre-treatment**<br>**(2 hours)** | **Treatment**<br>**(22 hours)** |
| --- | --- | --- |
| Control | Blank D3 media | Blank D3 media |
| TNF-alpha | Blank D3 media | TNF-alpha (20<br>ng/mL) |
| Dynasore | Dynasore (80 μM) | Blank D3 media |
| TNF-alpha +<br>Dynasore | Dynasore (80 μM) | TNF-alpha (20<br>ng/mL) |

### Statistical analysis

All statistical tests were conducted using GraphPad Prism (GraphPad Software, CA). Statistical significance of differences between controls and treatment groups was assessed using a two-tailed unpaired t-test or one-way ANOVA, depending on the experimental design.

## Results

### TNF alpha exposure enhances the brain uptake of Aβ

Following TNF-alpha infusion via the internal carotid artery in WT mice, the ^125^I-Aβ42 administered as bolus injection via the femoral vein (Figure 1A) exhibited a biexponential disposition in plasma (Figure 1B). The plasma pharmacokinetics of ^125^I- Aβ42 were unaltered in mice infused with TNF alpha compared to control mice (Figure 1B). However, the permeability of ^125^I-Aβ42 at the BBB in TNF alpha-infused mice, assessed as the PS product, was approximately two-fold (**p<0.01, *p < 0.05; two-way ANOA) greater in the cortex, hippocampus, and thalamus compared to control mice (Figure 1C). In addition, the whole brain accumulation of ^125^I-Aβ42 was assessed as the percent of injected ^125^I-Aβ42 radioactivity normalized to brain weight (%ID/g), which was approximately 2-fold (*p < 0.05; two-tailed *t*-test) higher in WT mice infused with TNF-alpha compared to control mice infused with PBS (Figure 1D). Based on the Gjedde-Patlak graphical analysis, the brain influx (Ki) of ^125^I-Aβ42 was approximately 1.5-fold higher (*p < 0.05, two-tailed *t*-test) in the TNF-alpha-infused mice compared to the PBS-infused mice (Figure 1E).

**Figure 1:**
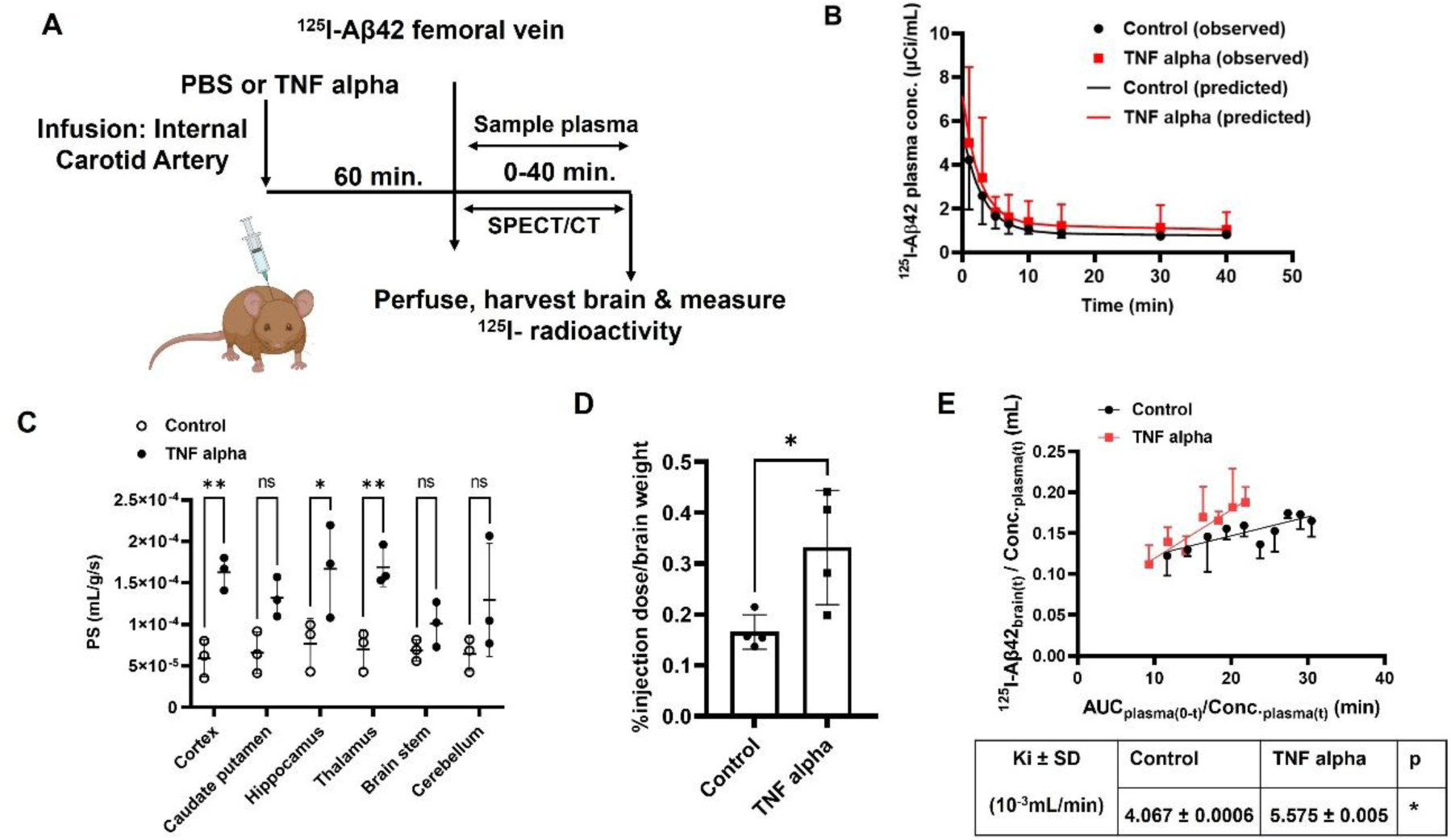
TNF alpha increases Aβ42 accumulation in wild-type mice following TNF alpha infusion. **(A)** Experimental scheme: Four-month-old wild-type female mice (B6SJLF1) were subjected to a pre-infusion of PBS or TNF-alpha via the left internal carotid artery over a duration of 15 minutes. Following the infusion, a waiting period of 60 minutes was observed before administering a bolus injection of ^125^I-Aβ42 (150 μCi) into the femoral vein. Blood samples were collected periodically from the femoral artery at intervals ranging from 0 to 40 minutes. These blood samples were subsequently centrifuged to isolate the plasma, and intact ^125^I-Aβ42 was precipitated using TCA. Radioactivity levels were quantified utilizing a gamma counter. After the final blood sampling event, the mice underwent transcardial perfusion with an excess of PBS, followed by dissection of brain regions. The level of ^125^I-Aβ42 activity within the brain was then assessed using a gamma counter. **(B)** Plasma concentration versus time profile of ^125^I-Aβ42 in mice infused with TNF alpha or PBS control. The observed values are presented alongside the predicted curves **(C)** The permeability-surface area (PS) products for ^125^I-Aβ42 for each brain region were estimated in mice infused with TNF alpha or PBS control. Data are expressed as mean ± standard deviation. Significance levels: *p < 0.05, **p < 0.01, two-way ANOVA with Bonferroni’s post-tests. **(D)** Bar graph depicts the percentage of the injected dose of ^125^I-Aβ42 in the whole brain per gram of tissue in TNF alpha-infused mice compared to PBS-infused mice Data are expressed as mean ± standard deviation. Significance levels: *p<0.05; determined by two-tailed t-test. **(E)** The brain influx (K_i_) of ^125^I-Aβ42 was estimated by the slope obtained from Gjedde-Patlak graphical analysis. Inset table values are mean ± SD (n = 4). Significance levels: *p < 0.05; two-tailed t-test.

### Increased uptake of Aβ42 by polarized BBB endothelial cell monolayers following TNF alpha stimulation

The impact of TNF-alpha stimulation on FITC-Aβ42 uptake by endothelial cells was investigated in PBMECs and hCMEC/D3 monolayers (Figure 2A). Upon TNF-alpha stimulation, the median fluorescence intensity (MFI), indicative of FITC-Aβ42 intracellular uptake, was approximately 2-fold (*p < 0.05, two-tailed *t*-test) higher in PBMECs (Figure 2B) and approximately 1.5-fold (*p < 0.05, two-tailed *t*-test) higher in hCMEC/D3 monolayers (Figure 2C) compared to control cells, as assessed using flow cytometry. The uptake of FITC-Aβ42 was further verified using confocal microscopy, which demonstrated increased FITC-Aβ42 fluorescence in hCMEC/D3 monolayers treated with TNF-alpha in comparison to control cells (Figure 2D).

**Figure 2.**
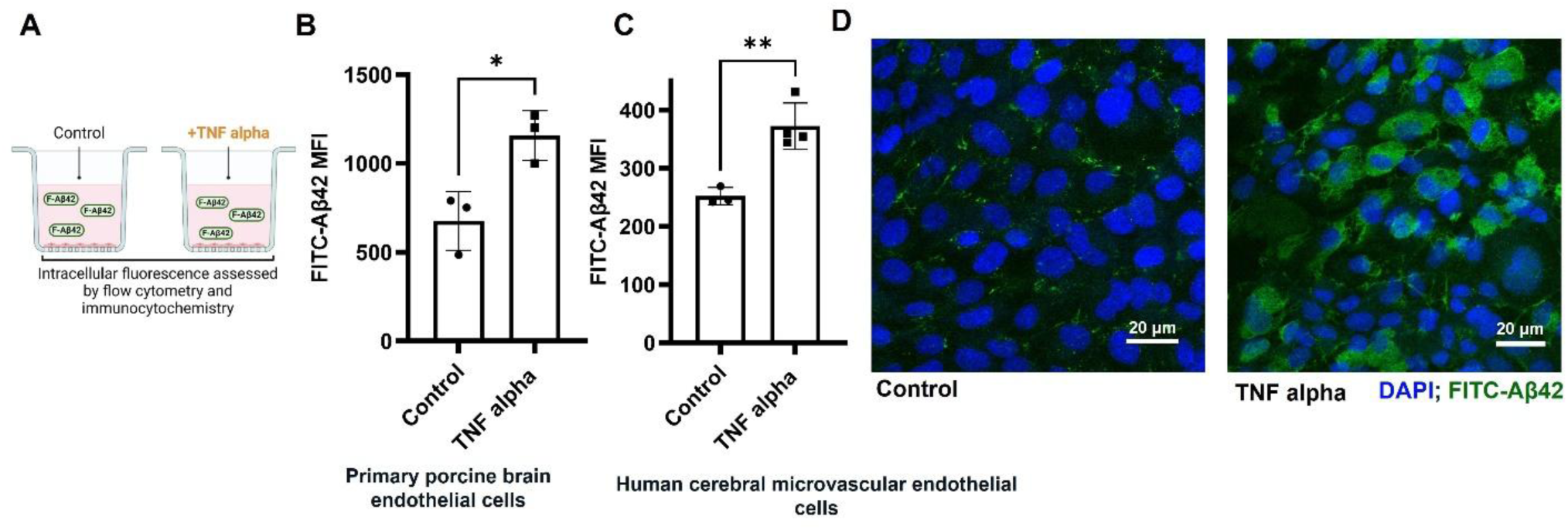
TNF-alpha increases FITC-Aβ42 accumulation in brain endothelial cells. **(A)** Experimental design. PBMECs or hCMEC/D3 cultured in 6 well plates were treated with TNF-alpha PBMECs (10 ng/mL), hCMEC/D3 (20 ng/mL) for 24 hrs. in low serum medium at 37°C. Subsequently, the cells were treated with FITC-Aβ42 (1 μM) and incubated for 1 hr. at 37°C. Intracellular fluorescence of samples was assessed by flow cytometry. **(B&C).** Bar charts display the intracellular median fluorescence intensity (MFI) of (B) PBMECs or (C) hCMEC/D3 incubated with FITC-Aβ42 in TNF-alpha treated and control groups. Data are expressed as mean ± S.D. Significance levels: *p<0.05, **p<0.01; determined by two-tailed t-test. **(D)** Confocal micrographs illustrate increased FITC-Aβ42 uptake in hCMEC/D3 cells treated with TNF-alpha compared to control cells. FITC-Aβ42 is visualized in green, while the nuclei are stained blue (DAPI). Scale bar = 20 μm

### Increase in FITC-Aβ42 uptake by BBB endothelial monolayers upon exposure to TNF alpha is mediated via downregulation of cofilin phosphorylation

Phospho-cofilin (serine 3) is an actin-binding proteins that regulate actin cytoskeleton remodeling(Kommaddi et al. 2018; Xu et al. 2021). We assessed their expression in polarized hCMEC/D3 monolayers following TNF-alpha stimulation using western blot analysis (Figure 4A). The cells treated with TNF-alpha demonstrated approximately 2 fold (p<0.05, two-tailed *t*-test) lower expression of phospho-cofilin compared to control cells (Figure 4B&D). Additionally, FITC-Aβ42 uptake in hCEMC/D3 monolayers was assessed by flow cytometry with or without siRNA-based cofilin knockdown followed by TNF-alpha stimulation (Figure 5A). The data demonstrates that the knockdown of cofilin increases endothelial uptake of FITC-Aβ42 by ∼1.5 fold (*p<0.05, two-tailed *t*-test) compared to control cells (Figure 5B). The efficiency of the cofilin knockdown was confirmed by western blots (Figure 5D&E).

**Figure 3:**
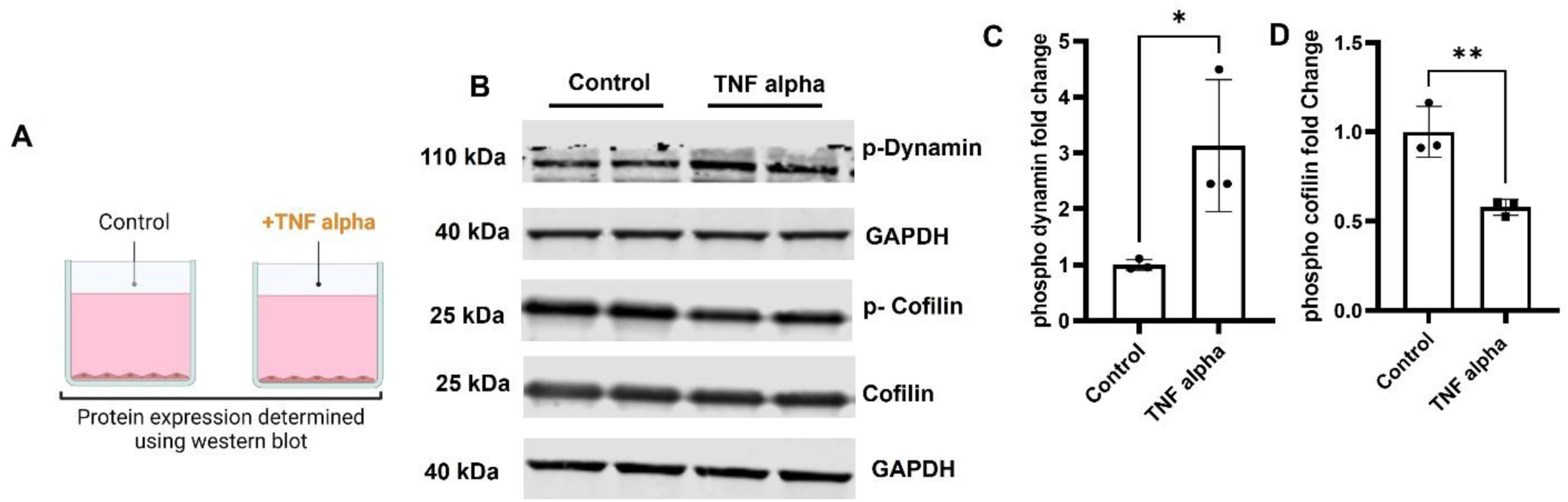
TNF alpha modulation of cofilin and dynamin phosphorylation. **(A)** Experimental design. hCMEC/D3 monolayers were cultured in 6 well plates and treated with TNF alpha (20 ng/mL) for 24 hrs. in low serum medium at 37°C. **(B)** Immunoblots depict phospho-cofilin, cofilin, phospho-dynamin, and GAPDH (loading control). **(C&D)** Bar chart shows quantification of immunoblots by densitometry for phospho-dynamin (C) and phospho-cofilin (D), both normalized to GAPDH protein levels. Data expressed as mean ± S.D. Statistical significance: *p<0.05, ***p<0.001; determined by two-tailed t-test.

**Figure 4:**
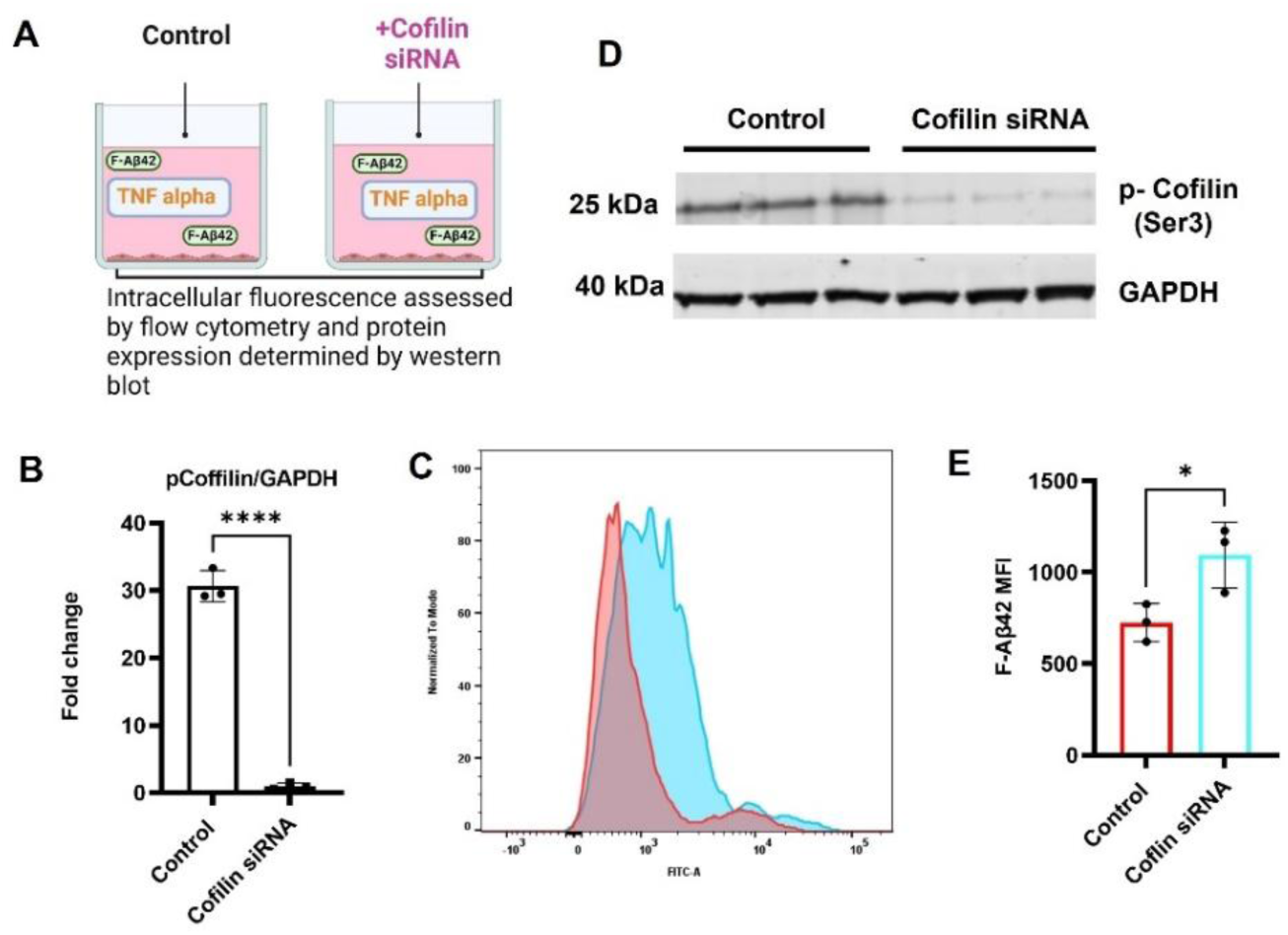
Impact of cofilin knockdown on FITC-Aβ42 uptake by brain endothelial cells. **(A)** Experiment design. hCMEC/D3 were transfected with or without cofilin siRNA (25 nM) for 6 hours. The treatment media was removed and replaced with blank D3 media for 24 hours at 37°C. Subsequently, the cells were treated with TNF alpha (20 ng/mL) in low serum (1%) medium at 37°C. The cells were then treated with FITC-Aβ42 (1 μM) and incubated for 1 hr. at 37°C. Intracellular fluorescence of samples was assessed by flow cytometry and protein expression was determined by western blot. **(B&C).** Histogram and bar chart depicting intracellular median fluorescence intensity (MFI) of hCMEC/D3 incubated with or without cofilin siRNA, in the presence of FITC- Aβ42 and TNF-alpha. Data expressed as mean ± S.D. Statistical significance: *p<0.05; determined by two-tailed t-test. **(D&E)** Bar chart demonstrates quantification of immunoblots by densitometry for phospho-cofilin, both normalized to GAPDH protein levels. Data expressed as mean ± S.D. Statistical significance: ****p<0.0001; determined by two-tailed t-test.

**Figure 5:**
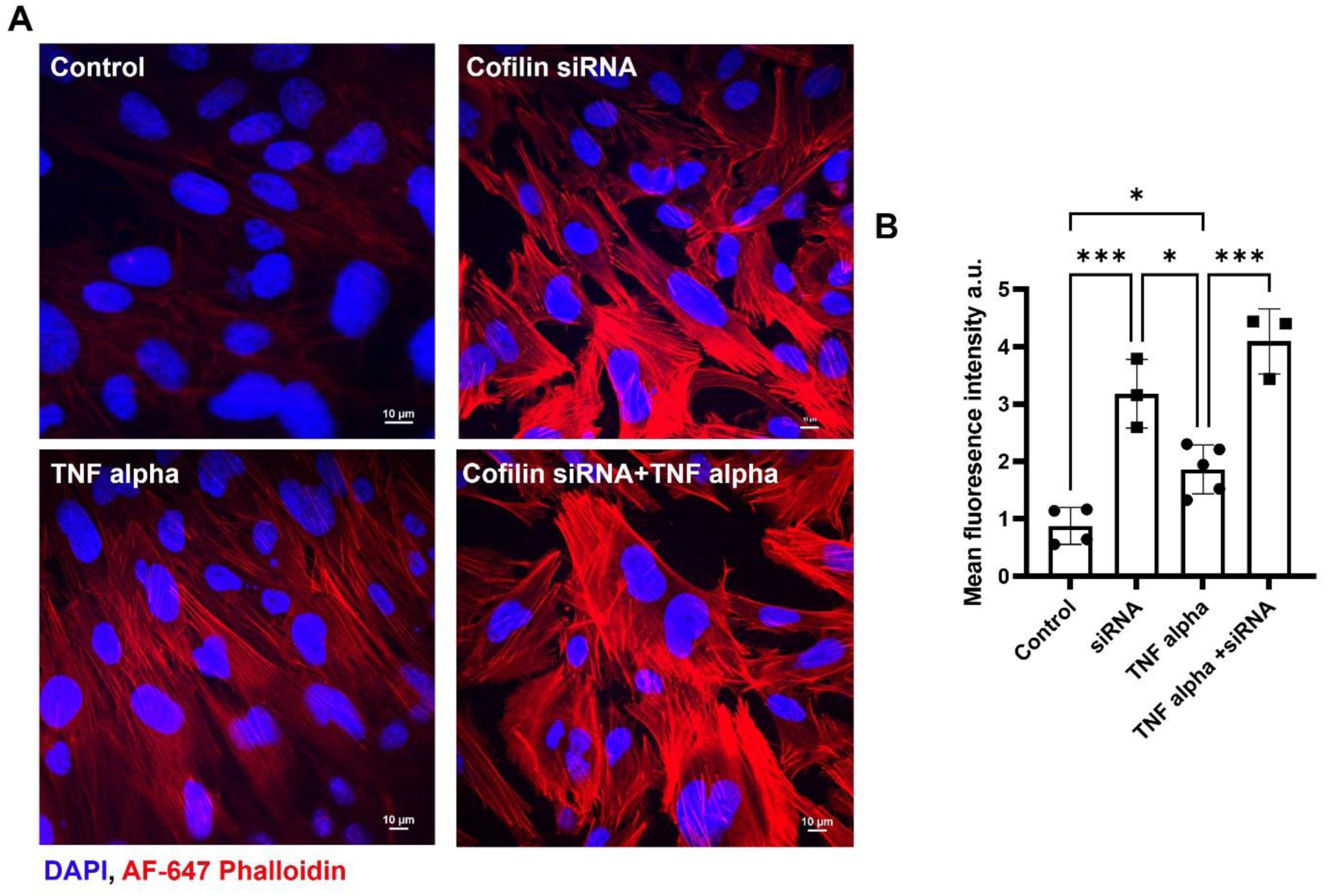
Impact of Cofilin Knockdown and Actin Inhibition on Cytoskeletal Dynamics in hCMEC/D3 Monolayers. **(A)** Fluorescence micrographs of AF-647 phalloidin in hCMEC/D3 monolayers. AF-647 phalloidin is visualized in red, while the nuclei are stained blue (DAPI). Scale bar = 20 μm **(B)** Bar char demonstrating mean fluorescence intensity of AF-647 phalloidin normalized to cell count in hCMEC/D3 monolayers. Data are expressed as mean ± SD *p < 0.05, **p < 0.01, two-way ANOVA with Bonferroni’s post-tests.

### Increased F-actin polymerization following cofilin knockdown

TNF alpha has been shown to enhance actin polymerization (Koukouritaki et al. 1999; Campos et al. 2009; Lim et al. 2016), which is significant as the organization of the actin cytoskeleton is essential for the invagination of endothelial membrane (Yang et al. 2022; Manenschijn et al. 2019). To evaluate the specific effects of TNF-alpha treatment and cofilin knockdown on actin polymerization, we investigated four distinct treatment groups: control, TNF-alpha, cofilin siRNA, and cofilin siRNA combined with TNF-alpha. We quantified actin polymerization in hCMEC/D3 monolayers by immunofluorescence microscopy of F-actin labeled with Alexa-Fluor 647 (AF-647) phalloidin. As shown in Figure 6A & B, TNF-alpha treatment alone significantly promotes actin polymerization compared to the control group. Similarly, cofilin knockdown enhances actin polymerization, as evidenced by the increased intensity of AF-647 phalloidin. The combined effect of cofilin knockdown and TNF-alpha treatment is additive, resulting in the highest level of actin polymerization among all treatment groups.

**Figure 6.**
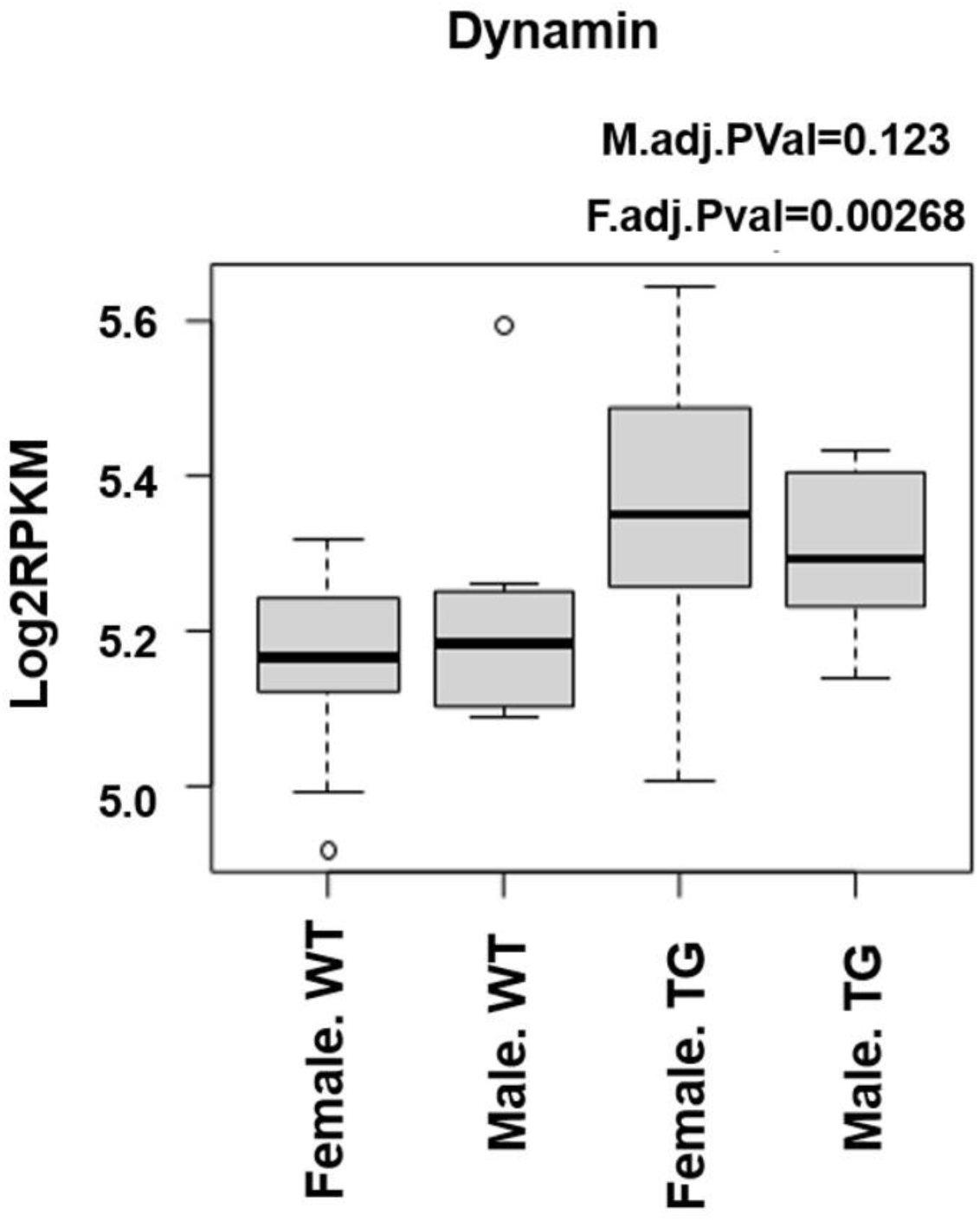
Increased dynamin expression in female AD transgenic (TG) mice. Box plot depicting higher dynamin gene expression in female APP/PS1 (TG) mice compared to female wild-type (WT) mice using the number of genes at a given reads per kilobases of transcript per 1 million mapped reads (RPKM). The data is plotted on a log2 scale.

### Increased dynamin mRNA expression in female AD transgenic (APP/PS1) mice

We investigated mRNA expression of dynamin, a GTPase protein involved in severing the endothelial membrane to release endocytic vesicles(Schmid and Frolov 2011), in WT and APP/PS1 mice using data extracted from a publicly available microarray dataset (GSE85162). The expression level in APP/PS1 was quantified based on the number of genes at a given RPKM, relative to age-matched WT controls. The dataset compared gene expression in male and female WT mice with male and female APP/PS1 mice, a well-established Alzheimer’s disease transgenic mice model that overexpresses Aβ42(Radde et al. 2006). A significantly elevated dynamin expression in female APP/PS1 mice compared to female WT mice (**p value<0.01). In contrast, no difference in dynamin expression was observed between male APP/PS1 and WT mice (Figure 7).

**Figure 7.**
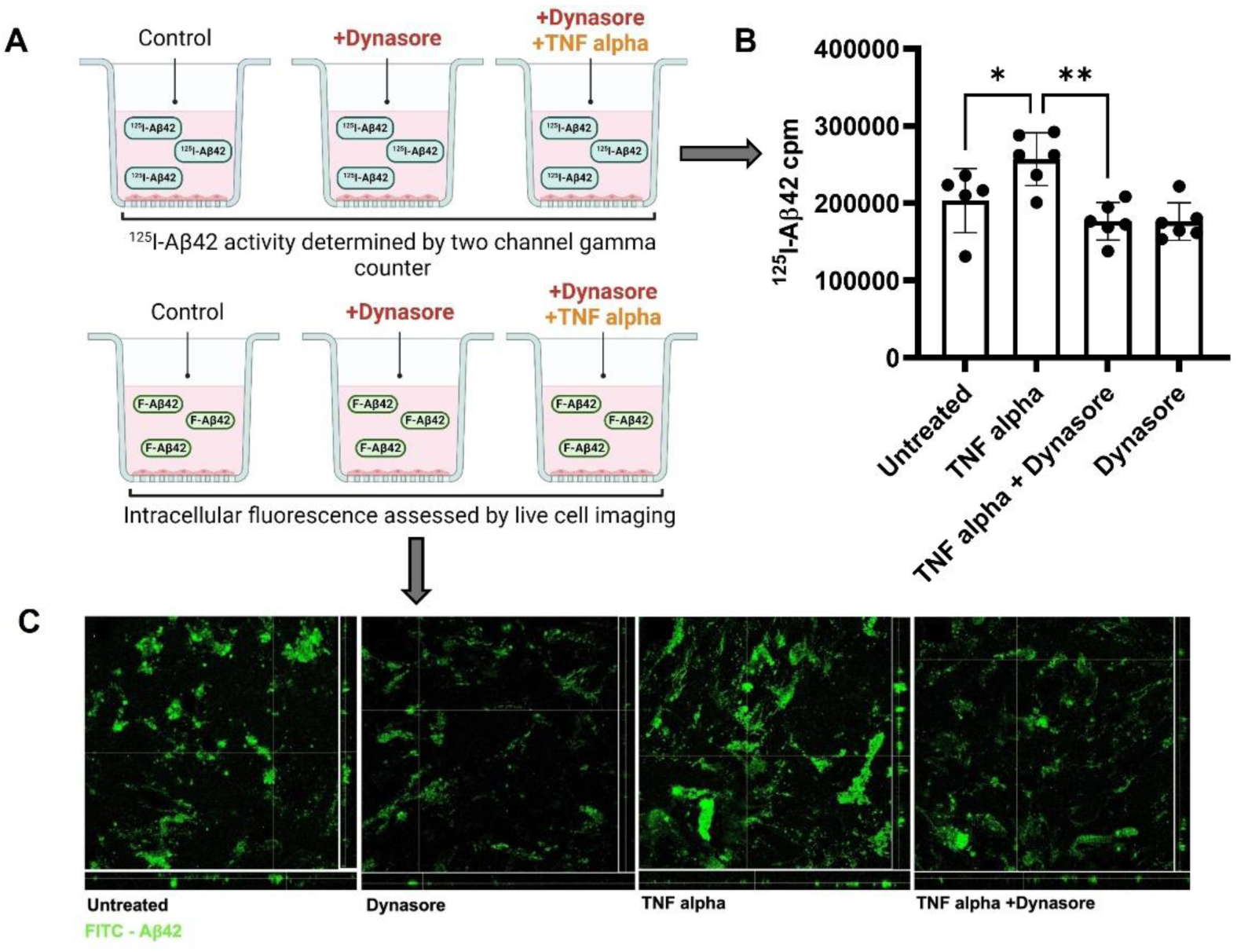
TNF alpha mediates Aβ42 brain endothelial cellular uptake via a dynamin-dependent mechanism. **(A)** Experimental scheme. hCMEC/D3 monolayers were pre-treated with dynasore (80 μM, 2hrs, 37 °C.) in low serum medium, followed by spiking the cells with TNF alpha (20 ng/mL) and incubating them for 22 hours at 37°C. Subsequently, the cells were treated with ^125^I-Aβ42 (5 μCi) or FITC-Aβ42 (1 μM) and incubated for 1 hr. at 37°C. Samples were assessed by measuring ^125^I activity using a gamma counter or by live cell imaging using confocal microscopy. (**B)** Bar chart demonstrates dynasore-mediated reduction in ^125^I-Aβ42 counts per minute (cpm) following treatment with TNF alpha. Data expressed as mean ± S.D. Statistical significance: *p<0.01, *p<0.05; determined by one-way ANOVA. (**C)** Dynasore inhibits TNF alpha induced increase in FITC-Aβ42 (green) endothelial cellular accumulation, as depicted in the Z-stack confocal micrographs. Scale bar (20 μm).

### Dynamin dependent uptake of Aβ42 in BBB endothelial cells stimulated with TNF alpha

To determine if TNF-alpha mediated increase in Aβ42 uptake is dynamin-dependent, the effect of TNF-alpha on phospho-dynamin expression was assessed by western blot. It was shown that hCMEC/D3 monolayers treated with TNF-alpha demonstrated a ∼3 fold (*p<0.05; two-tailed t-test) increase in phospho-dynamin levels compared to control cells (Figure 4A&C). Furthermore, we used dynasore, a chemical inhibitor of dynamin, to assess its influence on TNF-alpha-mediated increase in Aβ42 uptake in polarized hCMECD3 monolayers (Figure 8A). The hCMEC/D3 monolayers pretreated with dynasore, followed by treatment with TNF alpha and ^125^I- Aβ42 demonstrated ∼1.5-fold (**p<0.01, two-tailed *t*-test) reduction in ^125^I- Aβ42 uptake compared to cells treated with TNF-alpha alone (Figure 8B). Live cell imaging confirmed that TNF-alpha and dynasore-treated hCMEC/D3 monolayers exhibited reduced accumulation of FITC-Aβ42, as visualized through Z-stack images, compared to endothelial cells treated with TNF- alpha alone (Figure 8C).

**Figure 8:**
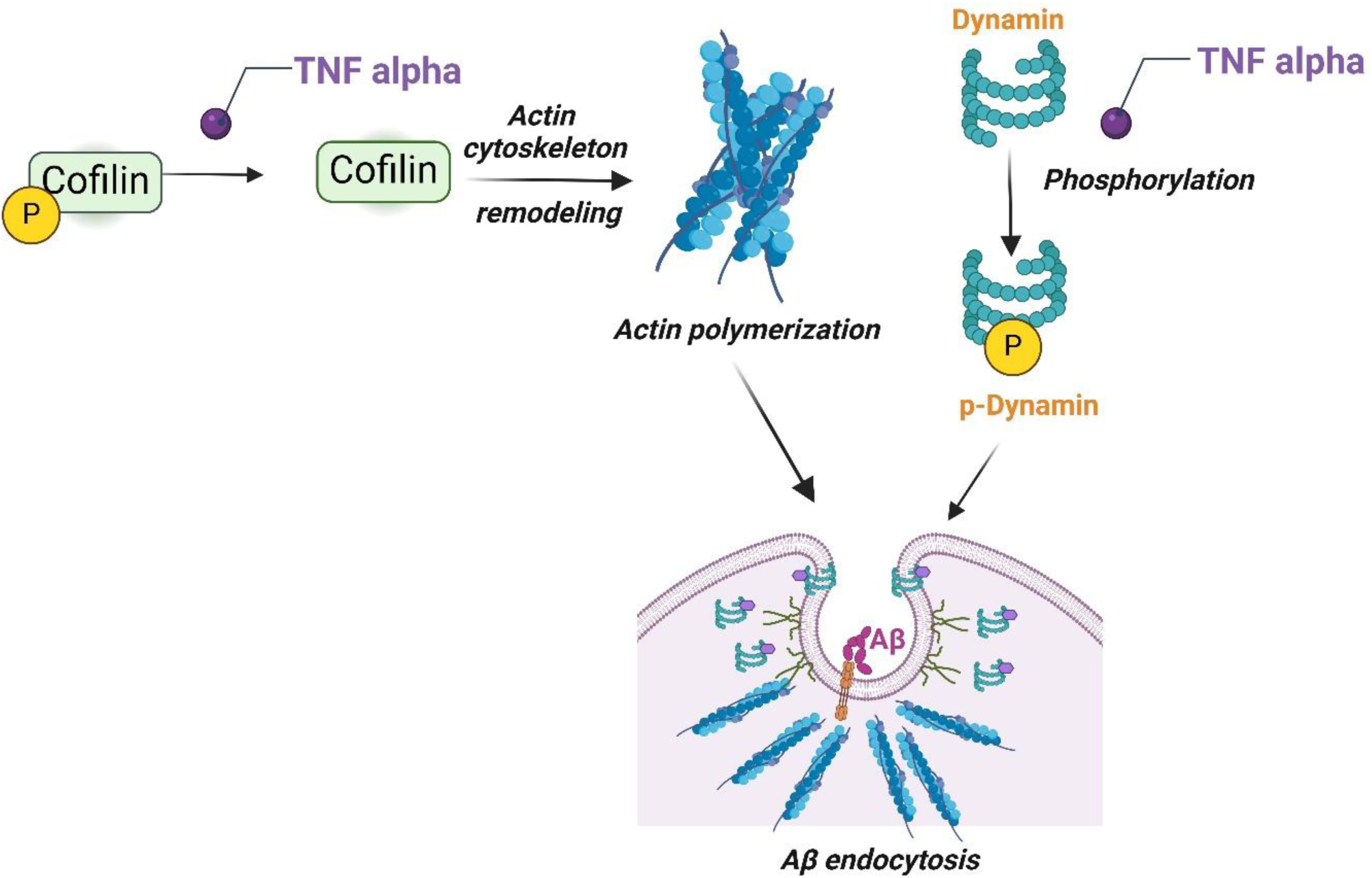
TNF alpha mediates an increase in Aβ42 endothelial cellular uptake via the cofilin-dynamin signaling pathway. TNF alpha induces a reduction in cofilin phosphorylation at the serine 3 site, facilitating actin cytoskeleton remodeling and actin filament polymerization. This process promotes endothelial membrane invagination, facilitating Aβ42 uptake by BBB endothelial cells. Additionally, TNF alpha enhances dynamin phosphorylation, which promotes membrane scission and endocytic vesicle release containing Aβ42.

### TNF alpha does not promote lysosomal degradation of Aβ42

Polarized hCMEC/D3 monolayers were co-incubated with FITC- Aβ42 and LysoTracker (marker for lysosomes) following treatment with TNF-alpha to examine lysosomal accumulation of FITC- Aβ42 (Figure 9A). The FITC- Aβ42 displayed no co-localization with lysosomes following TNF alpha treatment (Figure 9B).

**Figure 9.**
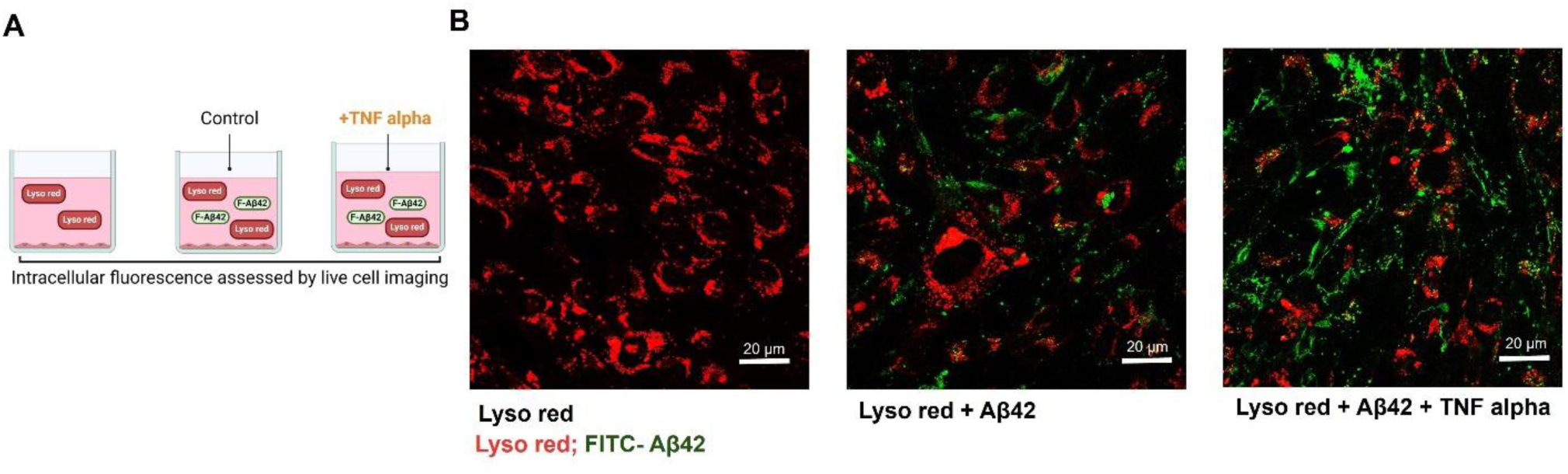
TNF alpha does not promote Aβ42 lysosomal degradation. **(A)** Experimental scheme: hCMEC/D3 monolayers cultured on 35 mm glass coverslip round bottom dishes were treated with TNF alpha (20 ng/mL) 24 hours in low serum medium at 37°C. Subsequently, the cells were incubated with FITC-Aβ42 (1 μM) for 30 minutes at 37°C, followed by co-incubation with LysoTracker red (75 nM) for another 30 minutes at 37°C. The treatment medium was then replaced with low serum D3 media (1% FBS), and live cell imaging was conducted using a confocal microscope. **(B)** Confocal micrographs illustrate the absence of colocalization between lysosomes and FITC-Aβ42 in the presence of TNF alpha. FITC-Aβ42 is visualized in green, while lysosomes are stained red. Scale bar = 20 μm

### Evaluating TNF Alpha Specificity in Aβ42 Uptake in BBB endothelial cells Using Insulin as a peptide Control

To determine if the effect of TNF alpha on Aβ42 uptake is specific or mediated by a general endocytic mechanism, we examined the influence of TNF alpha on insulin uptake in blood-brain barrier endothelial cells. The insulin uptake by hCMEC/D3 monolayers decreased by ∼1.8 fold (***p<0.001, two-tailed *t*-test) following treatment with TNF-alpha (Figure 10A&B). Furthermore, immunoblots of hCMEC/D3 cell monolayers treated with TNF-alpha showed a 3 fold (**p<0.01, two-tailed *t*-test) increase in phospho-IRS expression (serine 307) (Figure 10C&D), which impairs the ability of IRS-1 to associate with the insulin receptor and endocytose insulin (Paz et al. 1997).

**Figure 10:**
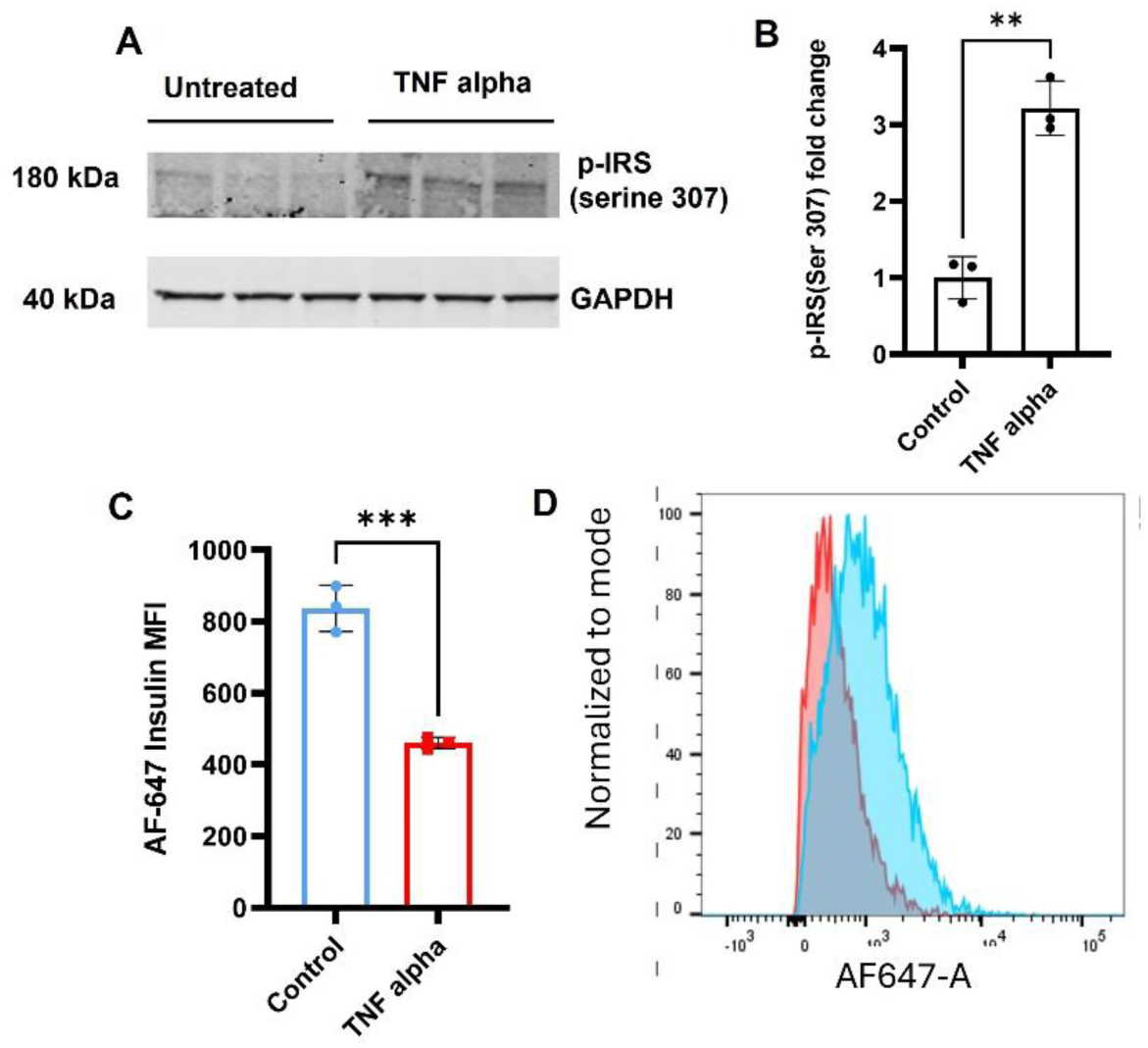
TNF alpha decreases AF647-insulin uptake in polarized hCMEC/d3 monolayers. Experimental design. hCMEC/D3 monolayers cultured in 6 well plates were treated with TNF alpha (20 ng/mL) for 24 hrs. in low serum medium at 37°C. Subsequently, the cells were treated with or without AF647-insulin (100 nM) and incubated for 20 minutes at 37°C. Intracellular fluorescence of samples was assessed by flow cytometry, and protein expression was determined by western blot. **(A)** Immunoblots depict phospho-IRS and GAPDH (loading control) **(B)** Quantification by densitometry **(C&D)** Bar chart and histograms depict median intracellular fluorescence (MFI) of AF-647 labeled insulin in TNF-alpha treated and control group. Data: mean ± S.D. ***p<0.001, *p<0.05; two-tailed t-test.

## Discussion

It is widely accepted that Aβ peptides enhance the secretion of pro-inflammatory cytokines and activate downstream signaling pathways that drive blood-brain barrier (BBB) dysfunction. TNF-alpha is one such cytokine, with higher levels reported in the plasma of Alzheimer’s disease patients compared to healthy controls(Zuliani et al. 2007). TNF-alpha’s ability to disrupt the BBB integrity and its influence on increasing Aβ production from amyloid precursor protein has been well studied(Cheng et al. 2018; Gao and Bayraktutan 2023; Sibson et al. 2002). Furthermore, its influence on receptors such as RAGE and LRP-1 that traffic Aβ from blood to brain and brain to blood respectively have been well documented(Tanaka et al. 2000; Hsu et al. 2021; Kitazawa, Hsu, and Medeiros 2016). TNF-alpha circulating in the blood does not enter the mouse brain after IV administration(Beutler, Milsark, and Cerami 1985). Therefore, in the systemic circulation TNF-alpha likely triggers cerebrovascular inflammation at the BBB by acting on the endothelial cells and increasing the expression of vascular inflammatory markers such as vascular cell adhesion molecule-1 (O’Carroll et al. 2015). However, the influence of TNF-alpha on BBB endothelial dysfunction and its influence on the BBB endothelial endocytosis machinery, which regulates Aβ trafficking in Alzheimer’s disease, remains unknown. In this study, we investigated the role of TNF-alpha in altering the endocytosis of Aβ in BBB endothelial cells. We investigated molecular mediators that are critical for regulating the cellular mechanisms of TNF alpha mediated endocytosis. These include F- actin, a structural protein essential for cytoskeletal integrity; phospho-cofilin, which regulates actin dynamics; and phospho-dynamin, a GTPase involved in membrane vesicle scission,

Previous studies have shown that the administration of an endotoxin such as LPS, which inflames the BBB endothelium, increase Aβ42 influx from plasma to the brain. However, the effect of inflammatory cytokines in the systemic circulation on Aβ42 influx at the BBB is not well understood(Jaeger et al. 2009). In this study, we investigated the impact of TNF-alpha, a pro-inflammatory cytokine, by infusing it via the internal carotid artery in WT mice on Aβ42 influx from plasma to the brain. Several studies have used this TNF alpha infusion model to examine the influence of an inflammatory stimulus on biological processes such as neuroinflammation, behavior, and locomotor activity (Biesmans et al. 2015; Skelly et al. 2013; Wang, Ando, and Dunn 1997). The permeability of Aβ42 at the BBB upon TNF-alpha exposure was assessed by determining the permeability surface area product (PS), a widely used parameter for assessing the uptake of macromolecules at the BBB. Following TNF-alpha exposure, Aβ42 PS values demonstrated an increase in the brain regions characterized by significant Alzheimer’s-related pathology, including the cortex, hippocampus, and thalamus. (Figure 1C). In addition, the Gjedde-Patlak plot generated from SPECT imaging provided an estimate of the plasma-to-brain transfer rate constant (Ki). The Ki of ^125^I-Aβ42 was significantly greater in TNF-alpha infused mice compared to control mice (Figure 1E).

The Aβ42 is trafficked from the plasma-to-brain via endothelial cells at the BBB. It is also important to investigate endothelial accumulation of Aβ42 in BBB endothelial cells. Published research studies have demonstrated that Aβ42 accumulation in endothelial cells drives BBB dysfunction by promoting insulin resistance, disrupting glucose transport, and stimulating the inflammation signaling cascade in the brain(Gali et al. 2019; Yamazaki et al. 2019; Stock, Kasus-Jacobi, and Pereira 2018). Hence, we evaluated the accumulation of FITC-Aβ42 peptide in BBB endothelial cell lines to investigate the molecular mechanisms driving TNF-alpha mediated increase in Aβ42 accumulation in the BBB endothelium. Using flow cytometry and confocal microscopy we demonstrated that TNF-alpha treated endothelial cells demonstrated a significantly higher intracellular accumulation of FITC-Aβ42 compared to control cells (Figure 2B-D). This data indicates that TNF-alpha does cause an increase in Aβ42 accumulation in BBB endothelial cells and potentially drives BBB dysfunction. Therefore, it is crucial to understand the molecular mechanisms underlying increased accumulation of Aβ42 in BBB endothelial cells upon TNF alpha exposure.

Wang et al. have demonstrated that Aβ42 endocytosis in BBB endothelial cells is energy, dynamin, and actin dependent(Wang et al. 2023). However, it is not clear how elevated TNF-alpha levels, impacts this endocytosis machinery. In this study, we demonstrated that TNF-alpha increases Aβ42 endocytosis in BBB endothelial cells by modulating key components of the endocytosis machinery such as actin, cofilin, and dynamin. This process occurs in two stages: membrane invagination and coated pit formation. The dynamic interaction of cofilin and actin(Xu et al. 2021; Qualmann, Kessels, and Kelly 2000) drives membrane invagination, while dynamin(Yang and Cerione 1999; McNiven 1998; Perrais 2022) regulates vesicle detachment (Figure 8).

Actin-binding proteins and actin-depolymerizing agents such as cofilin, play a critical role in mediating actin polymerization and depolymerization, thereby regulating the actin filament dynamics. The phosphorylation and dephosphorylation status of cofilin governs the activity of cofilin, whereby phosphorylation of cofilin at serine 3 site promotes severing and depolymerization of actin filaments as well as stabilization of actin filaments, which prevents membrane invagination. However, native form of cofilin promotes actin polymerization(Mizuno 2013), which in turn, promotes membrane invagination that drives endocytosis(Yang et al. 2022; Manenschijn et al. 2019). Here, we demonstrated that TNF-alpha exposure to hCMEC/D3 monolayers decreases phospho-cofilin levels (Figure 4). Our studies have demonstrated that knocking out cofilin in hCMEC/D3 monolayers using cofilin siRNA followed by TNF-alpha exposure increased the uptake of FITC-Aβ42 (Figure 5) compared to control cells. Furthermore, an increase in AF-647 phalloidin staining intensity, which is indicative of an increase in actin polymerization that in turn promotes endothelial membrane invagination was higher in the TNF-alpha group and cofilin KO group compared to the control group. However, a combined treatment of cofilin KO and TNF-alpha had an additive effect and demonstrated a higher degree of actin polymerization in comparison to the TNF-alpha alone group. Studies have demonstrated that cofilin is vital for the regulation of actin cytoskeleton dynamics and is controlled by the phosphorylation of cofilin(Lappalainen and Drubin 1997; Gunsalus et al. 1995; Abe et al. 1996; Nebl, Meuer, and Samstag 1996). These results indicate that TNF alpha promotes endothelial membrane invagination, which in turn mediates Aβ endocytosis through its influence on the actin-cofilin network.

After the membrane invagination step governed by actin-cofilin signaling pathway, which allows for Aβ42 endocytosis, the next step is the detachment of the newly formed vesicle, which is governed by dynamin. Dynamin is a GTPase protein that is involved in membrane scission followed by vesicle release. Several papers have demonstrated that Aβ42 endocytosis is dependent on dynamin(Choi et al. 2019; Wesen et al. 2017; Cirrito et al. 2008; Wang et al. 2023). Our studies have further demonstrated that the dynamin mRNA gene expression levels in Alzheimer’s disease female transgenic mice were higher in comparison to control female mice. This could be correlated to amyloid burden since female APP/PS1 mice have higher brain amyloid burden in comparison to male mice(Wang et al. 2003). In addition, our studies concluded that treatment of hCMEC/D3 monolayers with TNF-alpha significantly increases phospho-dynamin levels (Figure 4), where activation of this phospho counterpart aids in membrane scission and release of the endocytic vesicle. Alternatively, pre-treatment of hCEMC/D3 monolayers with a small molecule inhibitor of dynamin such as dynasore, inhibits the GTPase activity of dynamin, preventing its phosphorylation, and consequently blocking vesicle formation(Macia et al. 2006). Our studies demonstrated that dynasore pre-treatment followed by stimulation with TNF alpha decreased Aβ42 uptake in comparison to TNF-alpha alone treated cells (Figure 8). Thus, TNF alpha mediated increase in Aβ42 uptake is most likely mediated by modulating phospho-dynamin levels.

Next, we assessed lysosomal accumulation of FITC-Aβ42 following TNF alpha treatment to investigate if TNF alpha can promote lysosomal degradation of Aβ42 once endocytosed. Our studies show that TNF alpha does not promote lysosomal accumulation of Aβ42 within one hour, as no colocalization between F- Aβ42 and LysoRed was noted (Figure 9). Further studies need to be conducted to understand the intracellular changes of Aβ42 accumulation in early, late, and recycling endosomes of endothelial cells following TNF alpha stimulation to assess its effects on the intraendothelial transit of Aβ42.

To verify if the TNF alpha’s effect on Aβ42 uptake is specific or mediated by general changes in the endocytotic machinery, we investigated the influence of TNF alpha on insulin uptake in BBB endothelial cells. Insulin and Aβ42 peptide share similarities, where they are both polypeptides that contain regions of β-sheets within its structure(Hardy and Selkoe 2002; Dodson and Steiner 1998) and have a hydrophobic core(Jarrett, Berger, and Lansbury 1993; Hua and Weiss 2004). Our studies have demonstrated that treatment of hCEMEC/D3 monolayers with TNF-alpha decreased AF- 647 insulin uptake compared to control cells (Figure 10 A&B). We also demonstrated that TNF-alpha treatment decreases insulin uptake by increasing phospho-IRS (serine 307) levels (Figure 10C&D), which diverts insulin signaling from its normal pathway of tyrosine autophosphorylation of the β subunit of insulin to an inhibited signaling state that could potentially disrupt insulin uptake(Rui et al. 2001; Hotamisligil 1999) by endothelial cells. Therefore, the TNF alpha-mediated mechanism for Aβ42 peptide uptake appears to be peptide-specific.

In summary, our study demonstrated that the presence of TNF-alpha in the systemic circulation in mice increases the influx of Aβ42 peptide into the brain. Furthermore, we showed that TNF-alpha promotes internalization of Aβ42 by the BBB endothelial cells. This endocytosis process is dependent on key components of the BBB endocytosis machinery involving cofilin-actin and dynamin dependent processes. These results provide a mechanistic interpretation of the influence of TNF-alpha in promoting uptake of Aβ42 in the BBB endothelium. Further research is needed to understand TNF- alpha’s effect on mechanisms involved in the intracellular transit of endocytosed Aβ42 in endothelial cells and their potential role in progression of BBB dysfunction in AD.

## Footnotes

This work was supported by the Minnesota Partnership for Biotechnology and Medical Genomics [Grant 00056030], National Institute of Health [Grant AG058081], and National Institutes of Health/National Institute of Neurological Disorders and Stroke [R01NS125437].

## Conflict of interest statement

No author has an actual or perceived conflict of interest with the contents of this article

## Acknowledgements

Figures were created on Biorender.com

## Data Availability Statement

The authors declare that all the data supporting the findings of this study are contained within the paper.

## Authorship contributions

*Participated in research design:* V.S.S., K.K.K, K.R.K., and V.J.L.

*Conducted experiments*: V.S.S. and G.L.C.

*Contributed to new reagents or analytic tools:* K.K.K, V.J.L., and K.R.K.

*Performed data analysis*: V.S.S. and X.T.,

*Wrote or contributed to the writing of the manuscript:* V.S.S., and K.K.K.

## Abbreviations

TNF alpha: Tumor necrosis factor alpha
BBB: Blood brain barrier
Aβ: Amyloid beta
WT: wild-type
RAGE: Receptor for advanced glycation end product
LRP-1: Lipoprotein receptor-related protein-1
hCMEC/D3: Human cerebral microvascular endothelial cell
PBMECs: Primary porcine brain microvascular endothelial cells
TG: Transgenic
FITC: fluorescein isothiocynate
DPBS: Dulbecco’s phosphate- buffered saline
TCA: Trichloroacetic acid
CytoA: cytochalasin A
TBST: Tris-buffered saline with 0.1% Tween20 detergent
FBS: Fetal bovine serum
MFI: Median fluorescence intensity

## Notes

### Competing Interest Statement

The authors have declared no competing interest.

### Summary of Updates

The version of the manuscript has been revised to update methods, figures and findings.

